# Environmental control of Pub1 (NEDD4 family E3 ligase) in *S. pombe* is regulated by TORC2 and Gsk3

**DOI:** 10.1101/2021.02.21.432158

**Authors:** Tingting Wang, Philip Woodman, Sean J. Humphrey, Janni Petersen

**Affiliations:** Flinders Health and Medical Research Institute, Flinders Centre for Innovation in Cancer, Flinders University, Adelaide, SA 5042, Australia; School of Biological Sciences, Faculty of Biology, Medicine and Health, Manchester Academic Health Science Centre, University of Manchester, Manchester, M13 9PT, UK; Charles Perkins Centre, School of Life and Environmental Sciences, The University of Sydney, Camperdown, NSW, Australia; Nutrition and Metabolism, South Australia Health and Medical Research Institute, North Terrace, Adelaide, SA 5000, Australia

**Keywords:** NEDD4, Pub1, TORC2, GSK3, Nutrient-stress

## Abstract

Cells respond to changing nutrient environments by adjusting the abundance of surface nutrient transporters and receptors. This can be achieved by modulating ubiquitin-dependent endocytosis, which in part is regulated by the NEDD4 family of E3 ligases. Here we report novel regulation of Pub1, a fission yeast *Schizosaccharomyces pombe* member of the NEDD4-family of E3 ligases. We show that nitrogen stress inhibits Pub1 function, thereby increasing the abundance of the amino acid transporter Aat1 at the plasma membrane and enhancing sensitivity to the toxic arginine analogue canavanine. We show that TOR complex 2 (TORC2) signalling negatively regulates Pub1, thus TORC2 mutants under nutrient stress have decreased Aat1 at the plasma membrane and are resistant to canavanine. Inhibition of TORC2 signalling increases Pub1 phosphorylation, and this is dependent on Gsk3 activity. Addition of the Tor inhibitor Torin1 increases phosphorylation of Pub1 at serine 199 (S199) by 2.5-fold, and Pub1 protein levels in S199A phospho-ablated mutants are reduced. S199 is conserved in NEDD4 and is located immediately upstream of a WW domain required for protein interaction. Together, we describe how the major TORC2 nutrient-sensing signalling network regulates environmental control of Pub1 to modulate the abundance of nutrient transporters.

## Introduction

In all eukaryotic cells the external environment regulates cell fate. Highly conserved TOR (Target Of Rapamycin) signalling plays a key role in this control by responding to environmental cues, including stress and nutritional availability. This is achieved through TOR control of a series of metabolic processes, cell growth, migration, division and differentiation. TOR signalling is extremely sensitive to changes in the cellular nutrient environment, and it is well established that reduced cellular energy levels and changes in amino acid concentrations are actively sensed to modulate TOR pathway activity (Laplante and Sabatini, 2012).

Several nutrient acquisition pathways support TOR control of anabolic cell growth (Selwan et al., 2016). These include autophagy, which breaks down cellular components to generate nutrients for anabolism, uptake of nutrients via surface transporters and receptor-mediated uptake of macromolecular nutrients (Kim and Guan, 2015; Laplante and Sabatini, 2012; MacGurn et al., 2011; Rispal et al., 2015; Roelants et al., 2017). Cells respond to alterations in their nutrient environment by regulating the abundance of surface nutrient transporters and receptors, in part by controlling their ubiquitin-dependent endocytosis.

Reciprocal regulation of TOR and nutrient pathways has been established, because nutrients activate TOR, whilst TOR activity promotes endocytosis and inhibits autophagy. The mechanisms of TOR’s inhibition of autophagy to promote rapid cell proliferation in high nutrient environments is well-established (Kim and Guan, 2015). However, the impact of TOR stimulated endocytosis on nutrient utilisation is complex. On the one hand, TOR-controlled enhancement of endocytosis removes ion, carbohydrate and amino acid transporters from the plasma membrane, and may also reduce the surface population of macromolecular nutrient receptors, altogether reducing nutrient uptake (Ghaddar et al., 2014; Piper et al., 2014). On the other hand, endocytosis is also vital for the uptake of macromolecular nutrients such as LDL (May et al., 2003).

TOR signalling is comprised of two structurally and functionally distinct multi-protein complexes. TOR kinases form TORC1 and TORC2 (TOR Complex 1 and 2), which are defined by unique subunits that are highly conserved across species. In mammalian cells the protein Raptor defines (mTORC1), while Rictor is exclusive to mTORC2 (Laplante and Sabatini, 2012). In the fission yeast *S. pombe* model, the focus of this study, Mip1 is the functional homolog of Raptor in TORC1, whilst Ste20 (Rictor homolog) defines TORC2 (Alvarez and Moreno, 2006; Hayashi et al., 2007; Matsuo et al., 2007) and Gad8, an ortholog of human AKT and SGK, is a well-established substrate of TORC2 (Du et al., 2016; Ikeda et al., 2008; Matsuo et al., 2003).

Studies in both yeast and mammalian cells have established mechanisms of TOR regulated endocytosis and have documented both TORC1 and TORC2 dependent regulation of specific endocytic cargo (Gaubitz et al., 2016; Grahammer et al., 2017; MacGurn et al., 2011; Rispal et al., 2015; Roelants et al., 2017). For example, in *S. cerevisiae*, amino acid permeases such as Can1 are down-regulated under nutrient-replete conditions by their ubiquitination by Rsp5 (a NEDD4 family E3 ligase), a process requiring the Rsp5 adaptor Art1 (MacGurn et al., 2011). This process is conserved in mammalian cells (Boase and Kumar, 2015; Erpapazoglou et al., 2014; Piper et al., 2014; Rosario et al., 2016). Ubiquitinated Can1 is then recognized by endocytic ubiquitin adaptors such as Ede1/Eps15 and Ent1/epsin and incorporated into clathrin-coated vesicles. TORC1 stimulates Can1 uptake pathway via the Npr1-dependent phosphorylation of Art1 and Can1, thereby enhancing Can1 ubiquitination (MacGurn et al., 2011). In mammalian neurons NEDD4 is a key regulator of neurite growth (Boase and Kumar, 2015) and studies have identified NEDD4-1 mRNA as a prominent target of mTORC1 regulated translation (Hsia et al., 2014). Human NEDD4-1 and NEDD4-2 (NEDD4L) also regulate ubiquitin-mediated autophagy, an mTORC1 controlled process (Chen et al., 2019; Lee et al., 2020). In fission yeast, the abundance of the Aat1 and Cat1 amino acid permeases on the plasma membrane increases after nitrogen starvation, and a contribution of Pub1 (NEDD4 family of E3 ligases), Tsc1/2 (Laplante and Sabatini, 2012) (an upstream inhibitor of TORC1) and TORC1 pathway to this localization has been demonstrated by several laboratories (Aspuria and Tamanoi, 2008; Karagiannis et al., 1999; Matsumoto et al., 2002; Nakase et al., 2012; Nakase et al., 2013; Nakashima et al., 2014; Nefsky and Beach, 1996; van Slegtenhorst et al., 2004).

Each member of the NEDD4 HECT E3 ubiquitin ligase family comprises an amino-terminal Ca2+-phospholipid binding domain (C2), WW domains for protein to protein interaction, and a carboxy-terminal HECT domain containing its ligase activity (Boase and Kumar, 2015; Huang et al., 2019; Manning and Kumar, 2018). In the absence of Ca2+ binding to the C2 domain, conformational changes auto-inhibits NEDD4, whereas phosphorylation of NEDD4 on S347 and S348 by CK1 leads to its ubiquitination and degradation (Boase and Kumar, 2015). NEDD4-2 can also exist in an inactive form, since AKT1- and SGK1-mediated phosphorylation of S342 and S428 promotes 14-3-3 binding to block NEDD4-2’s interaction with its substrates. In contrast, AMPK and JNK phosphorylation at the carboxy-terminus is required for its activation (Boase and Kumar, 2015).

NEDD4 is expressed in most mammalian tissues and regulates a number of key substrates. Therefore, not surprisingly, dysregulation of NEDD4 ligases gives rise to a variety of diseases including cancer, cystic fibrosis, respiratory distress, hypertension, kidney disease, nervous system dysregulation and epilepsy (Boase and Kumar, 2015; Manning and Kumar, 2018). In summary, NEDD4 ligase activity is regulated at multiple levels, including translation, phosphorylation, binding to accessory proteins and control of protein turnover. Consequently, the molecular mechanisms of its regulation are complex and are not fully understood. In this study we used the fission yeast model system to gain further insights into the mechanisms responsible for regulating the activity of this key E3 ligase. We show that nitrogen stress inhibits Pub1 function. TOR complex 2 (TORC2) and Gad8 (AKT) signalling negatively regulates Pub1 through their control of Gsk3 activity. Phosphorylation of Pub1 at serine 199 (a site conserved in NEDD4) is increased following TORC2/AKT inhibition and therefore Gsk3 activation. In summary, we show that the major TORC2 nutrient-sensing signaling network regulates Pub1 to modulate the abundance of nutrient transporters.

## Results

### TOR complex 2 (TORC2) negatively regulates Pub1

The cellular response to nutrient starvation is, in part, to increase the abundance of surface transporters to facilitate greater uptake of nutrients from the environment. In budding yeast, when nutrients are plentiful TORC1 inhibits Npr1 kinase to allow Rsp5 ubiquitin-dependent endocytosis of transporters (MacGurn et al., 2011). However, upon nutrient starvation when TORC1 activity is inhibited so too is ubiquitin-dependent endocytosis, leading to higher levels of transporters at the plasma membrane (MacGurn et al., 2011). We previously undertook a global quantitative fitness profiling study to identify genes whose loss altered cell fitness in response to nitrogen stress. Not surprisingly, deletion of Pub1 (a NEDD4-family E3 ligase and the homolog of budding yeast Rsp5) increased cell fitness in response to nutrient stress (Lie et al., 2018). This is presumably because cells were able to import higher levels of nutrients due to reduced ubiquitin-dependent endocytosis of nutrient transporters.

With the aim of increasing our understanding of how Pub1 itself is regulated by changes to the cellular nutrient environment we exposed wild type cells to nitrogen stress, by changing the nitrogen source from good to poor (here we changed from ammonia to proline - EMM2 to EMMP). This resulted in a 60% decrease in Pub1 protein levels (Figure 1A) (Supplementary Figure 1A demonstrates that the antibodies detect Pub1). Therefore, in response to nitrogen stress when ubiquitin-dependent endocytosis is inhibited, the Pub1 E3 ligase is down-regulated.

**Figure 1:**
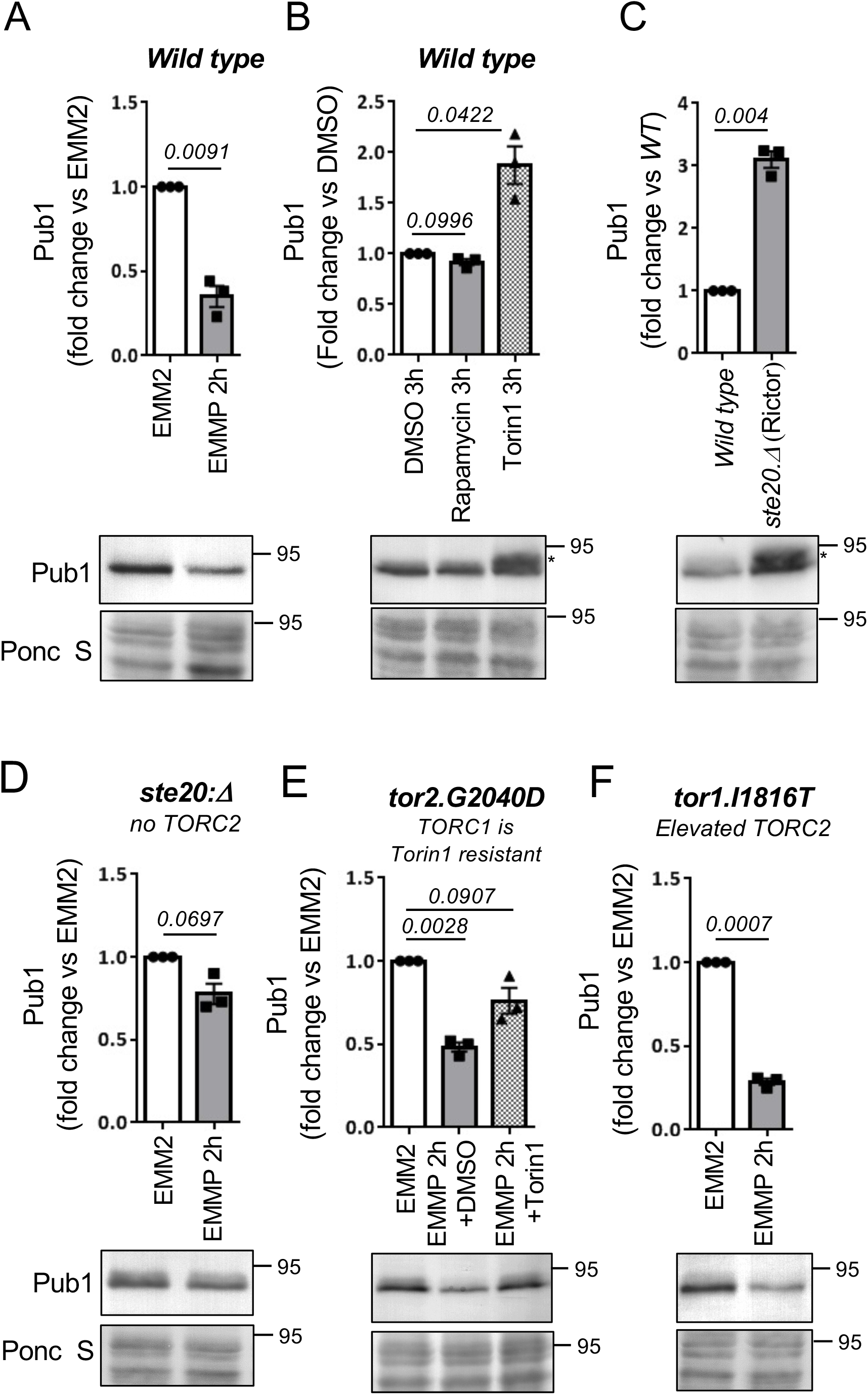
TOR complex 2 negatively regulates Pub1. Elevated Pub1 in cells lacking TORC2 function. A-F) Protein extracts were prepared from indicated yeast strains or treatments and immunoblotted for Pub1, Ponceau S is used to stain total protein. B-C) Following Torin1 treatment or inhibition of TORC2 a slower migrating form of Pub1 accumulates indicated by an asterisk E) Torin1 was added at 15 μM to EMMP medium of *tor2.G2040D* cells. Bars indicate fold change in levels vs. indicated controls ± s.e.m., *n* = 3, *n* represent biological independent experiments. representative immunoblots are shown.

To gain further insight into the environmental control of Pub1 protein levels, we treated wild-type cells grown in the good nitrogen source ammonia with the TOR kinase inhibitors rapamycin and Torin1 (ATP competitor TOR inhibitor 1 (Liu et al., 2012)) for 3hr. Rapamycin only inhibits TORC1 while Torin1 inhibits both TORC1 and TORC2 activities (Atkin et al., 2014). Rapamycin had no impact on Pub1 protein levels, whereas Torin1 promoted an increase in Pub1 levels (Figure 1B). At first glance this result appears contradicting. However, the imposition of nitrogen-stress to inhibit TORC1 has the opposite impact on TORC2 signalling, as previous reports demonstrated that TORC2 signalling is activated by nitrogen-stress in both fission yeast (after one hr of nitrogen withdrawal) and human cells (Hatano et al., 2015; Kazyken et al., 2019; Laboucarie et al., 2017; Martin et al., 2017). Since rapamycin had no impact on Pub1 levels, TORC1 is unlikely to have a major role in regulating Pub1 levels. Our data therefore suggest that it is the inhibition of TORC2 by Torin1 that results in increased Pub1 protein levels (Figure 1B), whereas, upon nitrogen-stress when TORC2 signalling is activated Pub1 levels decrease (Figure 1A). In agreement with this notion, deletion of the TORC2 specific component ste20 (Rictor) also increased levels of Pub1 (Figure 1C).

The impact that nitrogen-stress has on Pub1 protein levels was strongly diminished relative to wild-type when blocking TORC2 signalling in *ste20. Δ* (Rictor) mutants (Figure 1A, D). This indicates that active TORC2 is required for the observed decrease in Pub1 protein levels after nitrogen-stress. To test this further, we took advantage of our mutant in which we can inhibit TORC2 without affecting TORC1. Fission yeast Tor2 is the main kinase in TORC1, and we previously identified the *tor2.G2040D* mutation, in which TORC1 is resistant to Torin1 (Atkin et al., 2014). When the *tor2.G2040D* mutant was nitrogen stressed and Torin1 was added simultaneously (to inhibit only TORC2) the reduction of Pub1 due to media change to proline was diminished (Figure 1E). In contrast, in the TORC2 mutants Tor1.I1816T (Halova et al., 2013), which has a small increase in TORC2 activity, Pub1 levels were reduced more efficiently upon nitrogen-stress (Figure 1A, 1F).

Together, our data suggest that TORC2 negatively regulates Pub1 and that environmental control of Pub1 protein levels after nitrogen stress is regulated by elevated TORC2 signalling. Note that following Torin1 treatment a slower migrating form of Pub1 accumulates (indicated by an asterisk), indicating that TOR inhibition facilitates additional modification(s) of Pub1 (Figure 1C).

### Aat1 amino acid transporter localization to the plasma membrane upon nitrogen stress requires TORC2 activity

In fission yeast it is well-established that cells lacking Pub1 activity show increased abundance of the amino acid transporter Aat1 at the plasma membrane at cell tips (Matsumoto et al., 2002; Nakase et al., 2012). To visualize this, wild-type and *pub1::ura4*^+^ deletion cells were grown in EMM2, and wild-type cells were stained for 45 min with FM-4-64, which accumulates in the vacuoles, to differentiate between the two cell types when mixed 1:1 just before being imaged for Aat1.GFP localization (Figure 2A). Wild-type cells mainly had punctate cytoplasmic staining, previously attributed to localization at the Golgi (Liu et al., 2015). As expected, deletion of Pub1 increased Aat1 levels on the plasma membrane of cell ends by more than two-fold, relative to background fluorescence seen in the nuclei (Figure 2A), without changing overall levels of aat1.GFP protein (Supplementary Figure 1A). The same increase in Aat1.GFP localization in *pub1* deletion strains was seen when the *pub1.Δ* cells were stained for 45 min with FM-4-64 before mixing with wild type cells (Supplementary Figure 1B). Thus FM-4-64 does not interfere with Aat1.GFP localization or fluorescence.

**Figure 2:**
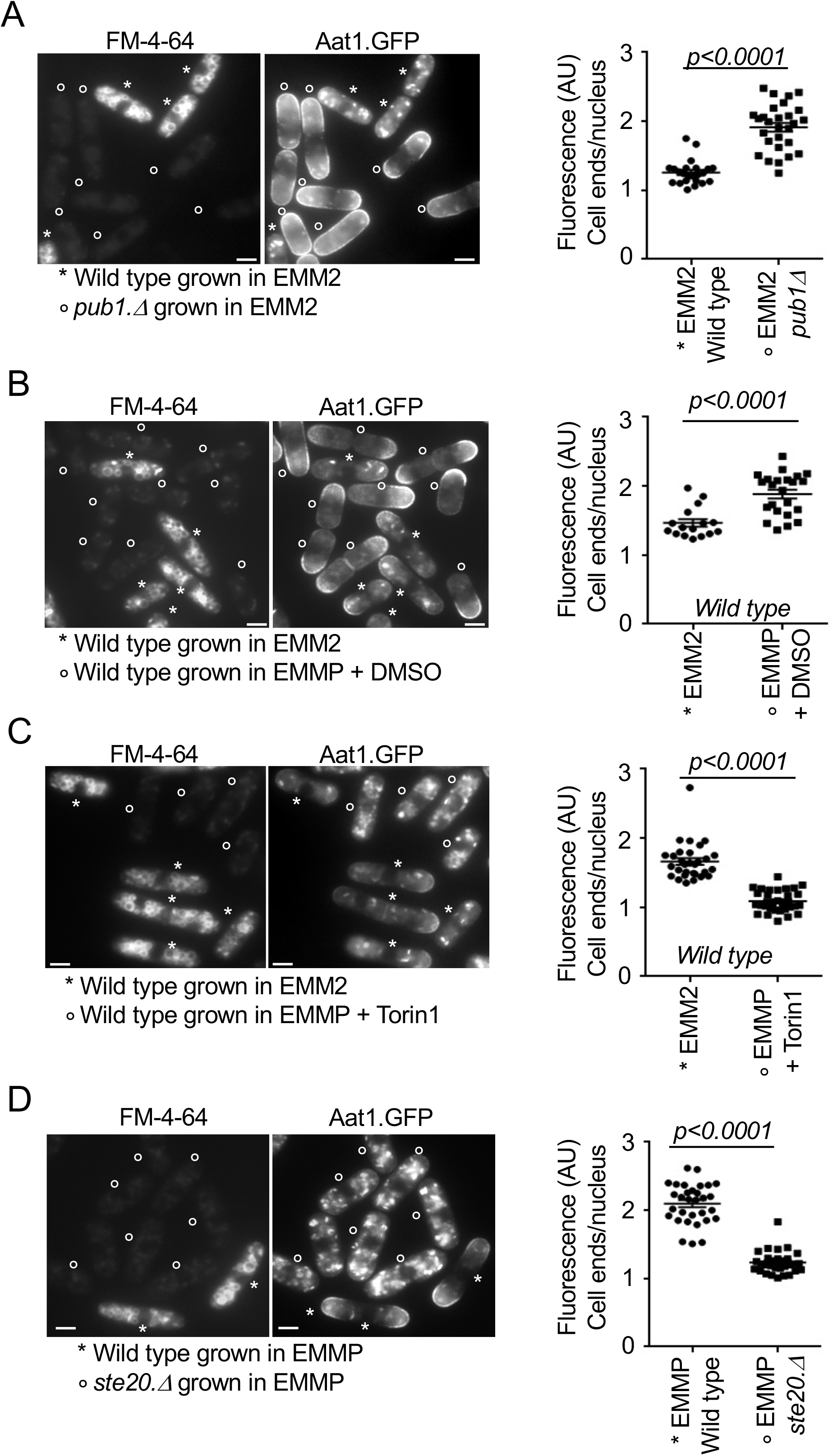
Aat1 amino acid transporter localization to the plasma membrane upon nitrogen stress requires TORC2 activity. A-D) Live cell imaging of Aat1.GFP in indicated strains. To differentiate between the two cell types or treatments (which were mixed at a ratio 1:1 immediately prior to imaging) control cells were initially stained with FM-4-64 (accumulates in the vacuoles, indicated by a star) for 45 min, before they were mixed with unstained cells (indicated by circles) for immediate imaging. The relative fluorescence intensity of Aat1.GFP in all cells was quantified as: intensity at cell ends/cell tips (where cell growth occurs) relative to nuclear background fluorescence levels in the same cell (value = 1 represent identical arbitrary fluorescence intensity at the growing cell tips and in the nucleus of the same cell). FM-4-64 staining does not affect Aat1.GFP fluorescence (supplementary Figure 1B) Scale bar = 3 μm. All stats are calculated from images from one experiment. Representative images are shown. Similar results were obtained for three independent biological repeats. A) Wild type cells were strained with FM-4-64 (indicated by a star), to differentiate between the two cell types when mixed 1:1 with unstained *pub1* deletion cells (indicated by a circle). B-C) Wild type cells grown in EMM2 were stained with FM-4-64. Unstained cells (indicated by a circle) grown in EMM2 were treated prior to media shift for 1 hour with DMSO (B) Torin1 at 25 μM (C). The cells were filtered into poor EMMP media supplemented with Torin1 or DMSO respectively for a further 90 min before live cell imaging together with stained cells. D) 90 minutes before imaging, wild type and *ste20* deletion cells were shifted from EMM2 to EMMP medium. The wild type cells were also stained with FM-4-64 (indicated by a star).

Imposition of nitrogen stress, by shifting wild type cells from EMM2 into EMMP reduced Pub1 protein levels by 60% (Figure 1A). Consistent with such a reduction in Pub1, localization of Aat1.GFP at the plasma membrane was increased in nitrogen stressed cells (Figure 2B), whilst total aat1.GFP protein levels remained unchanged (Supplementary Figure 1C). However, the addition of Torin1 upon nitrogen stress to inhibit TORC2 signalling (Atkin et al., 2014) and increase Pub1 protein levels (Figure 1B, C) abolished Aat1.GFP localization to the plasma membrane (Liu et al., 2015) (Figure 2C). Finally, nitrogen-stress of the *ste20* (Rictor) deletion, which block TORC2 function and thus increase Pub1 levels (Figure 1C) also blocked Aat1.GFP localization at the plasma membrane (Figure 2D). In summary, Aat1.GFP localization at the plasma membrane in poor nitrogen environments correlates with TORC2 regulation of Pub1 levels.

### TORC2 and Gad8 (AKT/SGK) are required for Pub1 dependent nutrient uptake

TORC2 regulates Pub1 protein levels and the abundance of amino acid transporters on the cell membrane (Figure 1 and 2). To gain further insight into the molecular mechanism of TORC2 control of Pub1 function, we assessed the levels of Pub1 proteins in cells deleted of the only known substrate of TORC2 in fission yeast – the Gad8 kinase (AKT/SGK homolog) (Du et al., 2016; Ikeda et al., 2008; Matsuo et al., 2003). Like cells deleted for the TORC2 specific component ste20 (Rictor) (Figure 1C), elevated Pub1 levels were also observed in cells deleted of *gad8* (Figure 3A). In the TORC2 mutants Tor1.I1816T, which has a small increase in TORC2 activity (Halova et al., 2013), Pub1 levels appeared slightly reduced although this was not significant. Furthermore, blocking TORC2 signalling in *gad8.Δ* mutants only reduced Pub1 protein levels by ~20% (Figure 3B) relative to a 60% reduction in wild-type cells (Figure 1A), indicating that active Gad8 (AKT) is required to downregulate pub1 following nitrogen stress.

**Figure 3:**
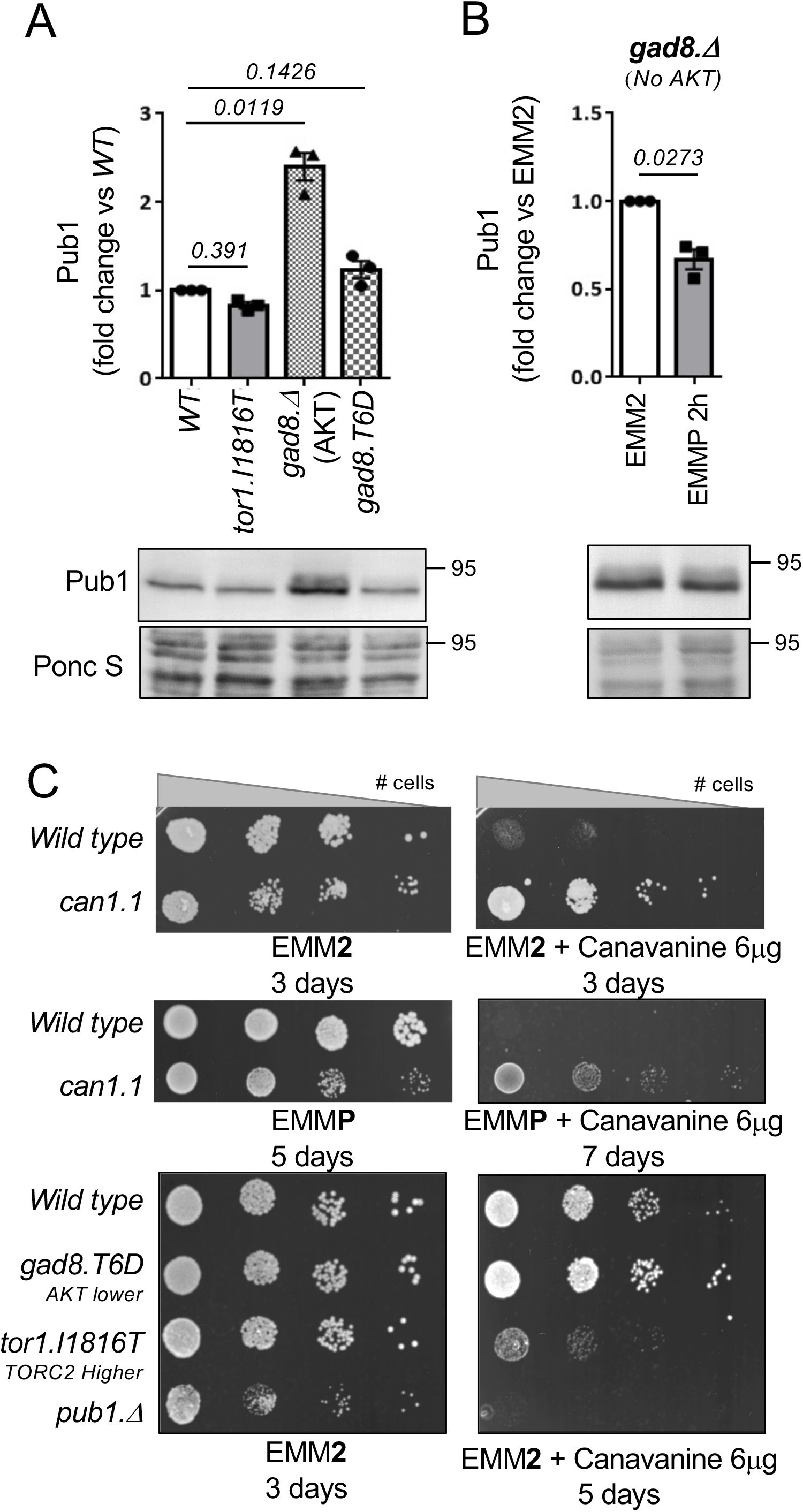
TORC2 and Gad8 (AKT/SGK) are required for Pub1 dependent nutrient uptake. A-B) Elevated Pub1 in mutants of Gad8 (AKT). Protein extracts were prepared from indicated yeast strains and immunoblotted for Pub1, Ponceau S is used to stain total protein. Bars indicate fold change in levels vs. indicated control ± s.e.m., *n* = 3. *n* represent biological independent experiments. C) Growth characteristics of indicated strains on EMM2 and EMM2 + 6 μg/ml Canavanine and on EMMP or EMMP + 6 μg/ml Canavanine. For all growth assays similar results were obtained for three independent biological repeats.

As shown above, cells increase the abundance of surface transporters to facilitate greater uptake of nutrients from the environment when stressed for nutrients (Figure 2B). We next used a simple well-established colony-forming growth assay to assess the role of TORC2 signalling on Pub1 function *in vivo*. Transport of canavanine into cells, a toxic arginine analogue, is in part regulated by the amino acid transporter Can1 (Fantes and Creanor, 1984), as the *can1.1* canavanine resistant mutant allele, in contrast to a wild-type strain, is able to form colonies when spotted from a serial dilution on agar-plates supplemented with canavanine. This is because the faulty transporter reduces the uptake of the toxic compound (Figure 3C) (Fantes and Creanor, 1984). In contrast, it is well-established that cells deleted for the Pub1 E3-ligase (which independently have reduced growth rate, even on EMM2 control media) are hypersensitive to canavanine (Figure 3C). This is because the block to ubiquitin-dependent endocytosis increases Can1 transporter abundance and therefore canavanine uptake (Aspuria and Tamanoi, 2008; Fantes and Creanor, 1984). Interestingly, *can1.1* resistance is reduced in poor nutrient environments (EMM2 vs EMMP) (Figure 3C), which is consistent with decreased Pub1 function in EMMP (Figure 1A) and therefore increased transporter levels on the plasma membrane. This finding suggests that additional transporters may transport canavanine in the absence of Can1 function in the *can1.1* mutant.

We next tested whether TORC2 and Gad8 control of Pub1 protein levels affected cells’ sensitivity to canavanine. Cells deleted of *ste20* and *gad8* display a substantially impaired growth rate compared to wild type cells (data not shown), so are not ideal candidates to assess growth rates in our “canavanine-sensitivity” assay. We therefore took advantage of two other mutant strains to assess the consequences of increased or decreased TORC2/Gad8 activity. We previously showed that whilst a Gad8.T6D mutant (which reduces Gad8 function, through reduced TORC2 binding to Gad8) has normal growth rates on EMM2 media (Figure 3C), Gad8 activity is reduced albeit not blocked (Du et al., 2016). Reduced Gad8 activity in Gad8.T6D cells resulted in somewhat larger colony size (increased cell proliferation) on canavanine plates when compared to wild-type cells (Figure 3C), indicating that Pub1 function was modestly increased in this mutant to reduce the uptake of toxic canavanine. Importantly, Pub1 levels were also slightly increased in Gad8.T6D cells (Figure 3B), consistent with the modest increase in growth rate on canavanine plates. Notably, the opposite impact on growth rates was observed in cells with enhanced TORC2 activity in the Tor1.I1816T mutant (Halova et al., 2013) (Figure 3C), as this mutant was sensitive to canavanine and exhibited a very slight reduction in Pub1 protein levels. Together, these observations indicate that TORC2 and its downstream substrate Gad8 negatively impact on Pub1 protein levels and therefore regulate the levels of transporters on the membrane, which can transport the toxic arginine analogue canavanine into cells.

### The TORC2 signalling pathway control Pub1 via Gsk3

To gain further insight into the molecular mechanism of TORC2 and Gad8 (AKT) control of Pub1 function we next considered Gsk3, as previous studies in human cells and fission yeast have shown that Gsk3 is a substrate of Gad8 (AKT) (Candiracci et al., 2019; Medina and Wandosell, 2011) and in fission yeast the TORC2 pathway was shown to regulate nutrient-dependent transcriptional elongation, through its inhibition of Gsk3 (Candiracci et al., 2019; Medina and Wandosell, 2011). Deletion of *gsk3* decreased levels of Pub1 by 50% relative to wild-type cells (Figure 4A). Fission yeast Gsk31 is an ortholog of Gsk3. Whilst Pub1 levels remained unaffected in the *gsk31.Δ* deletion strain, Pub1 protein levels were further reduced in *gsk3.Δ gsk31.Δ* double deletion when compared to *gsk3.Δ* (Figure 4A), indicating that the two Gsk3 kinases are capable of functional redundancy upon deletion (Miao et al., 2020; Qingyun et al., 2016). Although the growth rate of the *gsk3.Δ gsk31.Δ* double mutant on minimal EMM2 media is reduced (Figure 4B) it is sensitive to canavanine. Deletion of *gsk3.Δ* alone reduced colony size when exposed to the toxic compound, suggesting that Pub1 function was reduced in mutants lacking Gsk3 (Figure 4B). These observations fit with the reduced Pub1 protein levels seen in the *gsk3.Δ* mutant, which are further reduced in the *gsk3.Δ gsk31.Δ* double mutant (Figure 4A). We next asked whether the slow growth of *gsk3.Δ gsk31.Δ* double mutants (Figure 3B) could be rescued by Pub1 overexpression. Enhancing Pub1 levels only had a very minor enhancing impact on cell proliferation (Supplementary Figure 2A). This is in line with the numerous cellular functions of Gsk3 in cells (Xu et al., 2009).

**Figure 4:**
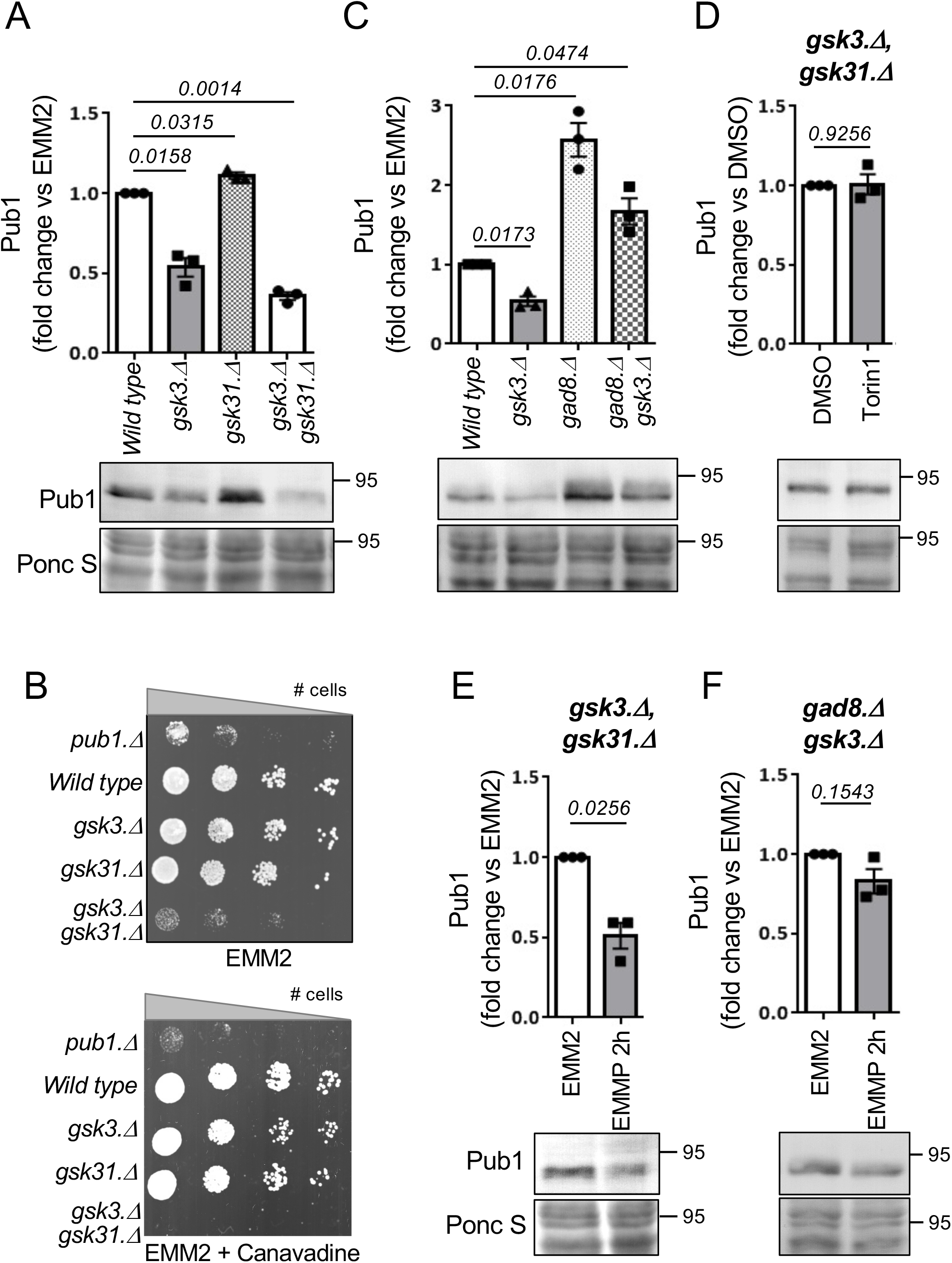
The TORC2 signalling pathway control Pub1 via Gsk3. Pub1 levels are reduced in mutants of Gsk3. A, C-F) Protein extracts were prepared from indicated yeast strains and immunoblotted for Pub1, Ponceau S is used to stain total protein. Bars indicate fold change in levels vs. indicated control ± s.e.m., *n* = 3. *n* represent biological independent experiments. Representative immunoblots are shown. B) Growth characteristics of indicated strains on EMM2 and EMM2 + 6 μg/ml Canavanine. For all growth assays similar results were obtained for three independent biological repeats.

We next analysed the reverse impact of Gad8 and Gsk3 on Pub1 levels further. If Gad8 regulates Pub1 through its demonstrated inhibition of Gsk3 activity, Pub1 levels in a double mutant are likely to resemble the levels seen in the *gsk3.Δ* mutant. The protein levels of Pub1 in the *gsk3.Δ gad8.Δ* double mutant were approximately half of those seen in *gad8.Δ* cells, and more than double that in *gsk3.Δ* cells (Figure 4C). These findings suggest that either the two kinases regulate Pub1 through independent mechanisms, or alternatively that the Gsk31 kinase present in the *gsk3.Δ gad8.Δ* double mutant is hyperactivated, due to lack of *gad8*, leading to elevated Pub1 protein levels compared to those seen in *gsk3.Δ*. Unfortunately, we were unsuccessful in generating a *gsk3.Δ gsk31.Δ gad8.Δ* triple deletion mutant to measure Pub1 protein levels and thus test this possibility. Nonetheless, in contrast to the situation in wild type cells (Figure 1B), chemically inhibiting TORC2 signalling and therefore Gad8 with Torin1 failed to increase Pub1 protein levels in the *gsk3.Δ gsk31.Δ* double mutant (Figure 4D). This observation implies that the increase in Pub1 protein levels seen in cells defective in TORC2 signalling (Figure 1B,C, 3A) is controlled by Gsk3 activation.

In response to nitrogen stress, the downregulation of Pub1 protein levels requires active TORC2 and Gad8 (Figure 1D, E 3B). Therefore, Gsk3 activity is predicted to be dispensable for this lowering of Pub1 protein levels after nitrogen-stress because increased TORC2 signalling under nitrogen stress would inhibit Gsk3 (Candiracci et al., 2019; Medina and Wandosell, 2011). Indeed, the reduction in Pub1 levels was maintained upon nitrogen-stress of single *gsk3.Δ* and *gsk31.Δ* mutants and the *gsk3.Δ gsk31. D* double mutant (Figure 4E, Supplementary Figure 2B). We next tested the possibility that Gsk3 overexpression blocks Pub1 downregulation following nitrogen stress. No significant change to Pub1 protein levels was seen in cells overexpressing Gsk3 compared to vector control (Supplementary Figure 3A). However, previous reports have failed to identify any strong phenotypes upon Gsk3 overexpression in fission yeast (Plyte et al., 1996; Qingyun et al., 2016) apart from rescue of cell growth in cells lacking AMPK activity at 37C, which was also observed here (Supplementary Figure 3B). Therefore, active Gad8 in nitrogen stressed cells may prevent significant activation of overexpressed Gsk3. We therefore turned to the *gsk3.Δ gad8.Δ* double mutant is which Gsk31 appears to be hyperactivated due to deletion of *gad8* deletion, leading to elevated Pub1 protein levels compared to those seen in *gsk3.Δ* (Figure 4C). Nitrogen stress of these cells, which lack the Gad8 inhibitor of Gsk31, failed to reduce Pub1 levels (Figure 4F).

In summary, our observations suggest that Gsk3 activity protects Pub1 and that the reverse impact of TORC2/Gad8 and Gsk3 on Pub1 levels comes about because of TORC2/AKT mediated Gsk3 inhibition (Candiracci et al., 2019) in fission yeast. Thus, lack of TORC2 activity enhances Gsk3 activity and consequently increases Pub1 protein levels.

### Gsk3 blocks Pub1 degradation by the proteasome

To further explore the mechanisms by which Gad8 and Gsk3 regulate Pub1 levels, we used quantitative PCR (qPCR) to assess the level of Pub1 mRNA in the two kinase deletion strains. Interestingly, whilst the Pub1 Protein levels are high in the *gad8.Δ* mutant, the mRNA levels are half that of wild type cells, whilst levels are unaffected in the *gsk3.Δ gsk31.Δ* mutants (Figure 5A). These observations suggest that the impact on protein levels in both mutants is independent of transcription. Auto-ubiquitination of *S. Cerevisiae* Rsp5 and SCF mediated degradation of human NEDD4 have been reported (Lam and Emili, 2013) (Liu et al., 2014). Interestingly, a block to proteasome function in the *mts3.1* proteasome mutant (Seeger et al., 1996) increased Pub1 levels three-fold compared to wild type, to reach levels similar to that seen in the TORC2 mutant (Figure 5B & 1C). We conclude that Pub1 is degraded by the proteasome. Blocking proteasome function rescued Pub1 protein levels in the *gsk3.Δ* mutant (Figure 5C), demonstrating that Gsk3 activity is essential to prevent Pub1 degradation by the proteasome. Furthermore, lack of proteasome function in the *mts3.1* mutant completely blocked Pub1 protein turnover (Figure 5D) following nitrogen stress, indicating that the proteasome is required for Pub1 destruction following under environmental nitrogen-stress.

**Figure 5:**
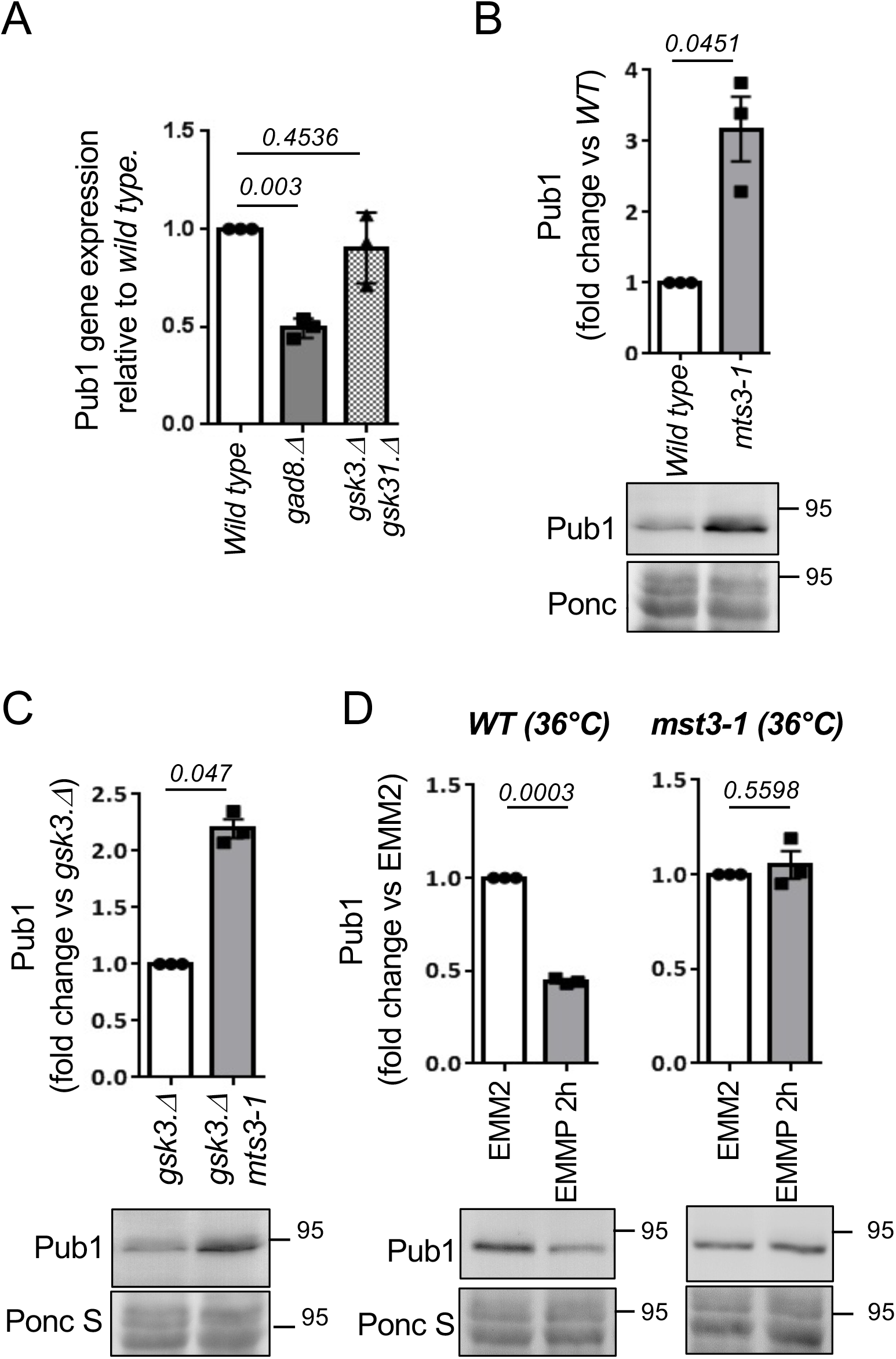
Gsk3 blocks Pub1 degradation by the proteasome. A) Levels of specific pub1 mRNA was measured by qPCR *n* = 3. Statistical significance was calculated Unpaired t test with Welch’s correction (Prizm). *n* represent biological independent experiments. B - D) Protein extracts were prepared from indicated yeast strains and immunoblotted for Pub1, Ponceau S is used to stain total protein. Bars indicate fold change in levels vs. indicated control ± s.e.m., *n* = 3. *n* represent biological independent experiments. Representative immunoblots are shown. D) Wild type and *mst3.1* cells were grown at 36 °C for 6 hours to inactive Mst3 before cell pelleting and protein extraction.

### Phosphorylation of Pub1 serine 199 enhances protein levels following TORC2 inhibition

To increase our understanding of how Gsk3 blocks proteasome-mediated Pub1 degradation we performed a quantitative, SILAC and label free mass spectrometry (MS) based analysis (Humphrey et al., 2018) to identify Pub1 phosphorylation. Protein extracted from wild type fission yeast that had been treated with Torin1 for two hours to inhibit TORC2 and therefore activate Gsk3 was mixed 1:1 with either SILAC labelled or label free solvent treated controls. This identified 5 phosphorylation sites on Pub1 (Supplementary table 1). Interestingly, Pub1 serine 199 (S199) phosphorylation, which was reported previously in global screens but not characterised further (Kettenbach et al., 2015; Swaffer et al., 2018), was upregulated 2.7-fold following Torin1 treatment. In contrast, upstream of S199, phosphorylation at serine 188 (S188) was decreased following Torin1 treatment (Supplementary Figure 4A, Supplementary table 1). S199 is located directly upstream of the first WW domain in Pub1 and both S188 and S199 are conserved in human NEDD4 upstream of WW domain 3 (Serine 824 and Serine 835; Figure 6A). Phosphorylation of human NEDD4 S824 and S835 have not been reported. However, trypsin which is used routinely in shotgun proteomics studies generates a relatively long (48 amino-acids) NEDD4 peptide including these sites, which may not be identified in the MS analysis. We managed to generate phospho-specific antibodies to Pub1 S188 (Supplementary Figure 4A,B). Consistent with our MS data, the relative levels of pS188 vs total Pub1 was reduced after 2 hr of Torin1 treatment (Supplementary Figure 4D). However, as pS188 is down regulated by Torin1, it is unlikely to represent a site phosphorylated by Gsk3.

**Figure 6:**
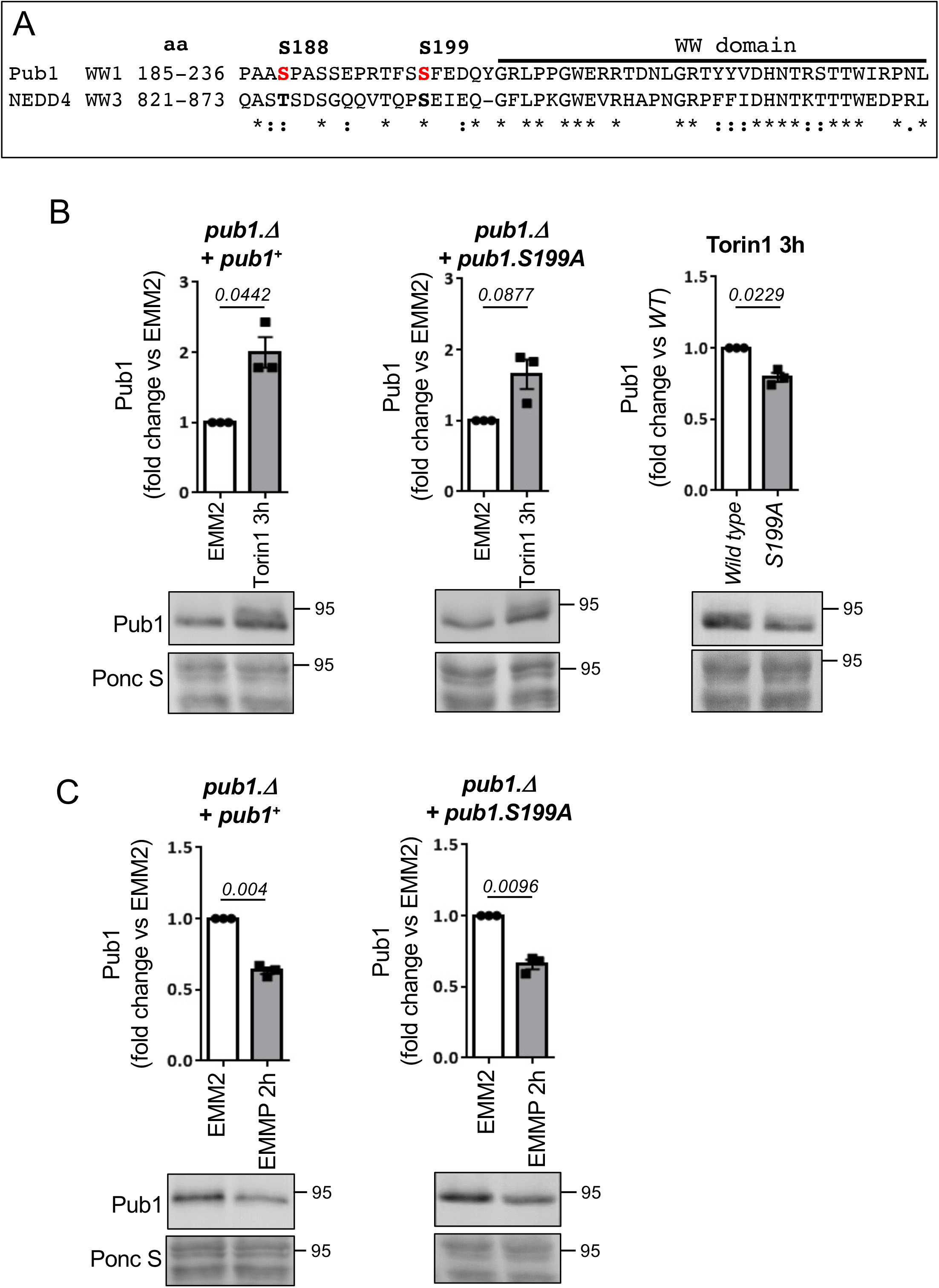
Phosphorylation of Pub1 serine 199 enhances protein levels following TORC2 inhibition. A) Sequence alignment of Pub1 S188 and S199 (shown in red) with human NEDD4 homolog, aa indicate amino acids. B, C) Protein extracts were prepared from indicated yeast strain and immunoblotted for Pub1, Ponceau S is used to stain total protein. Bars indicate fold change in levels vs. EMM2 ± s.e.m., *n* = 3. *n* represent biological independent experiments. Representative immunoblots are shown.

As described previously, in cells lacking Gsk3 activity Torin1 failed to increase Pub1 protein levels (Figure 4C) and failed to accumulate the slower migrating form of Pub1 (likely to represent the increase in phosphorylation) seen in wild type cells (Figure 1B). Gsk3 commonly phosphorylates a primed sequence S/T-X-X-X-S/T(P) pre-phosphorylated by another kinase (Beurel et al., 2015). However, priming-independent GSK3 phosphorylation has also been reported in cells, e.g. no priming kinase is required for LRP6 serine 1572 phosphorylation by Gsk3 (MacDonald et al., 2008; Zeng et al., 2008; Zeng et al., 2005). Interestingly, the sequence downstream of LRP6 S1572 is very similar to Pub1 S199 and NEDD4 835 (Supplementary Figure 4E), hence S199 may be a direct Gsk3 site. To analyse the role of Pub1 S199 phosphorylation following Torin1 treatment, we mutated the serine to a phospho-blocking mutant alanine (A). A *pub1* deletion strain was transformed with wild type Pub1 and the S199A mutant. Torin1 was able to enhance pub1 levels in both wild type and mutant S199A (Figure 6B), however Pub1 levels in Pub1 S199A failed to accumulate to the level of wildtype (Figure 6B).

Whether, Pub1 S199 is a direct Gsk3 site, remains to be established, however, it is unlikely to be the only site on Pub1 regulated by Gsk3, as Pub1 still accumulated in the S199A mutant albeit not to the level of wild type. Pub1 levels in unstressed condition was unaffected in S199A mutants (Supplementary Figure 4F). Furthermore, degradation following nitrogen stress was unaffected by the S199A mutation (Figure 6C), consistent with the notion that TORC2 signalling inhibits Gsk3 under nitrogen-stress, hence pub1 is still degraded in Gsk3 null cells (Figure 4E).

## Discussion

Here we show for the first time that the fission yeast NEDD4 family of E3 ligase Pub1 is regulated by the nutrient environment and the major nutrient sensing TORC2 pathway, to control the levels of amino acid transporter on the plasma membrane and thus nutrient uptake. Previous studies have established mechanisms of both TORC1 and TORC2 dependent regulation of specific endocytic cargo and membrane transport in both yeast and mammalian cells (Gaubitz et al., 2016; Grahammer et al., 2017; MacGurn et al., 2011; Rispal et al., 2015; Roelants et al., 2017). However, we now show that TORC2 and its downstream substrate Gad8 (AKT) negatively regulates Pub1 function via Gsk3 summarised in Figure 7. We demonstrate that Gsk3 protects Pub1 from proteasomal degradation, as blocking proteasomal function in the *mts3.1* mutant restores Pub1 protein levels in cells lacking Gsk3 activity (Figure 4D). Importantly, in both fission yeast and human cells, it is well established that AKT inhibits Gsk3 (Candiracci et al., 2019; Medina and Wandosell, 2011). Thus, the increase in Pub1 levels upon TOR inhibition with Torin1 is abolished in cells lacking Gsk3 activity (Figure 4D).

**Figure 7:**
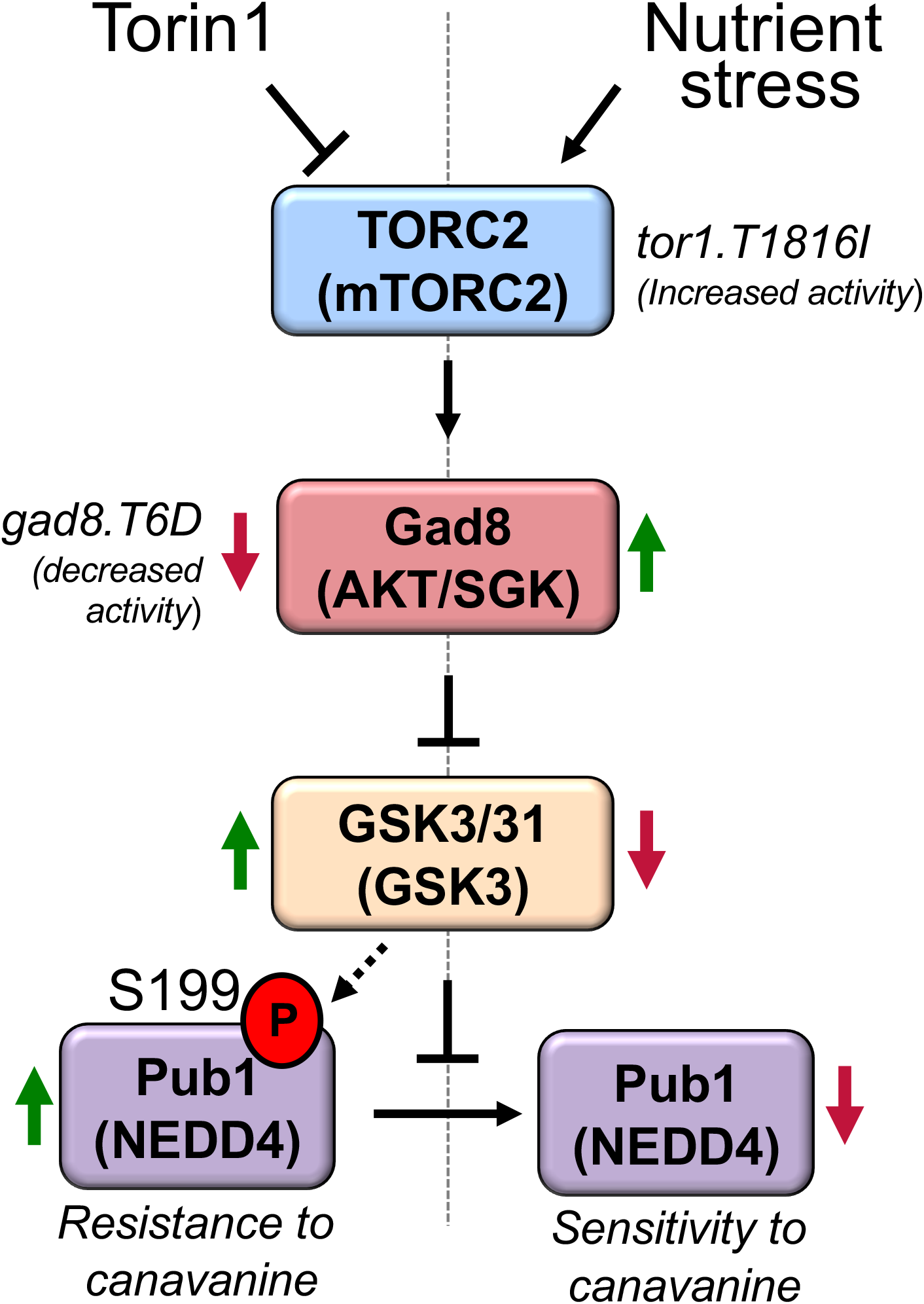
Schematic of TORC2 and Gsk3 dependent control of Pub1. TORC2 and Gad8 signalling regulates Pub1 through demonstrated inhibition of Gsk3 activity. Thus, reduced TORC2 activity (Torin1 or *gad8.T6D* mutant cells) enhances Gsk3 activity and consequently increases Pub1 Function. In contrast nitrogen stress known to activate TORC2 signalling (or *tor1.T1816I* mutant cells) inhibit Gsk3 activity to reduce Pub1 function.

When cells experience changes to their nutrient environment they respond by increasing the abundance of surface nutrient transporters, in part through down regulating their ubiquitin-dependent endocytosis. In agreement, we demonstrate TORC2-dependent Pub1 protein turnover through Gsk3 inhibition and proteasomal degradation following nitrogen stress (Figure 1D, 4F, 5D). This, in turn results in increased Aat1 amino acid transporter abundance on the plasma membrane (Figure 2B) and increased sensitivity to the toxic arginine analogue canavanine (Figure 3C) summarised in Figure 7.

Our previous investigation of global quantitative fitness to detect genes whose deletion altered cell fitness in response to nitrogen stress or inhibition of TOR signalling identified Pub1 (Lie et al., 2018). Deletion of *pub1* enhanced cell fitness, presumably because cells lacking *pub1* are able to import higher levels of nutrients, due to reduced ubiquitin-dependent endocytosis of nutrient transporters. In this screen the deletion of Gsk3 also enhanced the fitness of cells grown on minimal media (*p* = 0.108) (Lie et al., 2018), which is consistent with our observation that Pub1 protein levels are reduced in the *gsk3.Δ* mutant (Figure 4A). Increased viability upon nitrogen starvation of cells deleted of *gsk3* has also been reported in an independent genome wide screen (Sideri et al., 2015). A role for Gsk3 in protein stability is well-established, though in contrast to the protective role of Gsk3 on Pub1 protein stability we describe here, Gsk3 is known to prime many substrates for proteasome degradation, with over 25 substrates identified in human cells that are degraded in a Gsk3-dependent manner (Xu et al., 2009).

Whether Pub1 S199 is a direct substrate of Gsk3 remains to be established. Whilst Gsk3 commonly phosphorylates a primed sequence S/T-X-X-X-S/T(P) pre-phosphorylated by another kinase (Beurel et al., 2015) priming independent GSK3 phosphorylation has also been reported (MacDonald et al., 2008; Zeng et al., 2008; Zeng et al., 2005). The sequence downstream of Pub1 Serine 199 is similar to that of an established Gsk3 substrate (supplementary Figure 4E), highlighting this site as a candidate. Future experiments will establish whether Pub1 is a direct Gsk3 substrate. Human NEDD4 is also degraded by the proteasome, as phosphorylation of NEDD4 on S347 and S348 by CK1 leads to SCF mediated ubiquitination and degradation (Boase and Kumar, 2015). However, the SCF phospho degrons DSGXXS or T-P-P-X-S are not conserved in Pub1, and deletion of CK1 activity in fission yeast does not increase Pub1 levels (data not shown). How Gsk3 protects Pub1 from proteasomal degradation is currently unclear. However, Pub1 protein levels were reduced in the S199A mutant compared to wild type, when cells were treated with Torin1 (Figure 6B), suggesting phosphorylation of this site is important.

Pub1 S199 phosphorylation has no impact on protein levels in nutrient rich environments (Supplementary Figure 4F), therefore, considering the close proximity of Pub1 S199 and NEDD4 S835 to their WW domains (Figure 6A) phosphorylation is likely to regulate protein-protein interactions of adaptor proteins important for function under nutrient stress. Future experiments will address this.

Whether the regulation of Pub1 we report here is conserved in human cells is unclear at this stage. In human cells, GSK3 negatively regulates glucose homeostasis (Lee and Kim, 2007). Furthermore, insulin and growth factor signalling, which activates mTORC1, mTORC2, AKT and S6K inhibit GSK3 activity (Medina and Wandosell, 2011) and thereby increase glycogen synthesis. In contrast, NEDD4 enhances insulin and growth factor signalling (Cao et al., 2008; Fukushima et al., 2015). Our observation suggests that reduced GSK3 activity as a result of insulin signalling may decrease NEDD4 function and thus put a brake on insulin and growth factor signalling through a negative feedback loop. However, in human cells NEDD4 can directly bind to and ubiquitinate AKT which is prior phosphorylated on pS473, to degrade active AKT (Huang et al., 2020). Thus, decreased GSK3 activity and therefore reduced NEDD4 function would increase active AKT pS473, providing a positive feedback for insulin and growth factor signalling and glucose uptake to counteract the aforementioned negative feedback and thereby establish a steady state. Therefore, if conserved, the mechanism described here would most likely only impact hormone signalling and glucose uptake when this pathway is interacting with other signalling pathway(s) that alter the steady state.

In summary, here we show that Gsk3 protects Pub1 function, in part through S199 phosphorylation. We also provide the first evidence of NEDD4 family E3 ligase being regulated by nitrogen stress and TORC2 signalling to reduce ubiquitin-dependent endocytosis, thus increasing the abundance of amino acid transporters on the plasma membrane when nutrient levels are challenging.

## Acknowledgements

The Japanese national bio resource project for providing strains. This work was supported Australian Research Council [DP180101682] to SJH and JP, Flinders Foundation seeding grant and Flinders Universities supported this work. PW was supported by The Leverhulme Trust (RPG-2018-091).

The authors declare that they have no conflict of interest.

## Materials and Methods

**Table 1.**
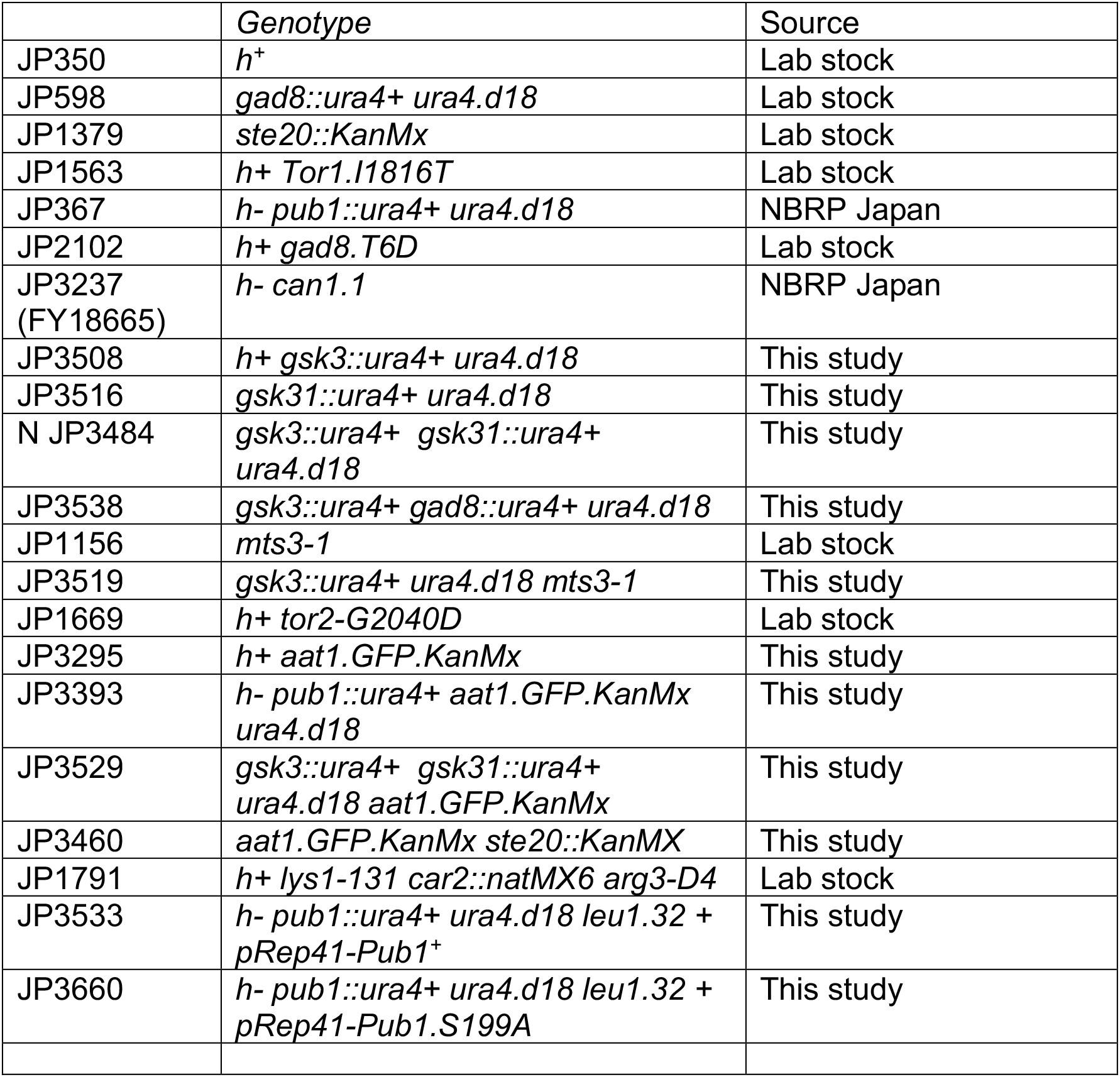
Strains used in this study

### Yeast cell cultures

All cultures were grown at 28°C and cultured in log phase for 48 h. Cells were inoculated in in Edinburgh minimal media (EMM2-N) (ForMedium) (Fantes, 1977) supplemented with NH_4_Cl (EMM2)(Petersen and Russell, 2016). Media change to (EMM2-N) (ForMedium) (Fantes, 1977) supplemented with proline (EMMP) (Petersen and Russell, 2016), was done by filtering cells, followed by resuspension in to prewarmed EMMP.

### SILAC labeling and harvesting culture for mass spectrometry

SILAC labeling: Cells were inoculated in Edinburgh minimal media (EMM2-N) (ForMedium) (Fantes, 1977) supplemented with 20 mM L-Glutamic acid (EMMG) (Petersen and Russell, 2016) and 75 mg/l of either light [L-arginine monohydrochloride (Sigma) and L-lysine monohydrochloride (Sigma)] or medium [lysine-L, 2HCl 4.4.5.5-D4 (Cat code DLM-2640, Eurisotop), arginine-L, HCl, U-13C6 99%13C (cat. no. CLM-2265, Eurisotop)] amino acids. Cells were cultured in log phase for 30 h to ensure complete incorporation of labelled amino acids into the proteome. Early log phase cultures at 3.5 × 10^6 cell/ml were treated with 15 **μ**MTorin1 or DMSO control. Cells were collected by filtration (MF-Millipore™ Filter, 1.2 μm pore size Cat. # RAWP04700, Millipore) washed with 15 ml TBS, resuspended in an appropriate volume of ice cold sterile ddH_2_O and dropped directly into liquid nitrogen to produce frozen cell droplets.

#### Mass spectrometry

SILAC mass spec analysis of samples processed using a SPEX Sample Prep LLC 6850 Freezer Mill in presence of liquid nitrogen, were performed as described previously (Humphrey et al., 2018). Data was analysed with MaxQuant (Cox and Mann, 2008) (v1.6.0.9) using the Andromeda search engine (Cox et al., 2011) to query a target-decoy database of *S. pombe* from UniProt (September 2019 release).

### Drug treatment

L-Canavanine sulfate salt (Cat. # C9758, SIGMA) was added to EMM2 agar plates at a concentration of 6 ug/ml, Torin1 (Cat. # 4247, TOCRIS) was used at a concentration of 15 μM and 25 μM. Rapamycin (Cat. #R0395, SIGMA) was used at a concentration of 300 ng/ml.

### Generation of *pub1.S199A* mutant

The *pub1* serine 199 point mutation was generated by site directed mutagenesis of Rep42-Pub1. Transformation of a *pub1::ura4+* deletion and selection on plates lacking leucine (Petersen and Russell, 2016) were used to select transformants. Strains were grown in the presence of 10 μM thiamine to reduce the level of Pub1 expression.

### Western blotting

TCA precipitation protocol was followed for *S.pombe* total protein extracts (Caspari et al., 2000). The following dilutions of antibodies were used in this study: 1/250 anti-Pub1 pS188 and 1/500 anti-Pub1 (custom made by Thermo Scientific, anti-Pub1 being non-specific to pS188) in PBS buffer, 1/500 anti-GFP (Cat. # 11814460001, Roche) in TBS buffer, skim milk was used as blocking agent. Alkaline phosphatase coupled secondary antibodies were used for all blots followed by direct detection with NBT/BCIP (VWR) substrates on PVDF membranes.

GraphPad Prism 6.07 was used for data analysis. Unpaired t test with Welch’s correction (Prizm version 7) were used for all Western blots. 95% confidence of interval was used for calculating significance.

### Fluorescent Microscopy

Staining of vacuoles: SynaptoRed™ C2 (Equivalent to FM®4-64) (Cat. # 70021, Biotium) was added to the growth media of cells (1 × 10^6 cells/ml) at a concentration of 1.5 μM for 45 min. Cultures of Stained and unstained cells were mix 1:1 and collected by filtration onto MF-Millipore™ Membrane Filter, 1.2 μm pore size (Cat. # RAWP04700, Millipore). Cells were resuspended in the original growth media of the FM®4-64 stained cells and subjected to live cell imaging immediately. Images of cells were obtained using a CoolSNAP HQ2 CCD camera. ImageJ were used to measure fluorescent intensities of Aat1.GFP. The relative fluorescence intensity (arbitrary units) of Aat1.GFP was quantified as: intensity at cell ends relative to nuclear background fluorescence signal of the same cell, to allow for comparisons of separate images and experiments. Statistical significance was calculated using Unpaired Students t test.

### RNA extraction and qPCR

RNA was extracted using TRIzol™ Reagent (Cat. # 15596026, ThermoFisher Scientific). In short, 1×10^7^ cells in early log phase were collected by centrifugation. Cell pellets were snap-frozen in liquid nitrogen. 1 ml of Trizol and 200 ul of glass beads (Cat. # 11079105, Biospec) were added to the cells. Cells were disrupted by a FastPrep-24™ (MP) at 5 m/s for 60 seconds for 3 cycles in a cold room. Cell lysate was processed according to manufacture’s instructions. RNA pellets were resuspended in 50 ul of RNAse free water. 1000 ng of RNA subjected to DNA digestion by TURBO DNA-free™ Kit (Cat. # AM1907).

First-strand cDNA were synthesized from 500 ng of RNA by using M-MLV Reverse Transcriptase, RNase H Minus, Point Mutant (Cat. # M3683, Promega). DNAse treated RNA, 500 ng Oligo (DT)_15_ (Cat. # C1101, Promega) and 100 ng random hexamer (Cat # C1181, Promega) and heated to 70 °C for 5 minutes, cooled to 4 °C, and incubated on ice for 5 minutes. For reverse transcription, RNA, primers, dNTP mix (Cat # N0446S, Bio New England Lab), M-MLV RT (H-) Point Mutant were used. 1: 4 diluted first-strand cDNA were used for second strand synthesis of cDNA and qPCR using Power SYBR™ Green PCR Master Mix (Cat. # 4367659, ThermoFisher Scientific). Reactions were run in Rotor-Gene Q (Qiagen) with initial activation at 95 for 10 minutes, followed by 40 cycles of 95 °C for 15 seconds, 58 °C for 1 minute. Comparative Quantitation Analysis from Rotor-Gene Q series software produced Representative Takeoff vale from triplicates of each sample. 2^−ΔΔCT^ method was used to calculate pub1 gene expression relative to housekeeping gene act1. Primers to amplify pub1 gene: Forward: CCCTTATTGGAATGAGACTTTTG; Reverse: GGGTCAACATTTCATCACCTC. Forward and Reverse Primers to amplify the control act1 gene was a described (Biswas et al., 2016).

---

**Supplementary Figure 1:**
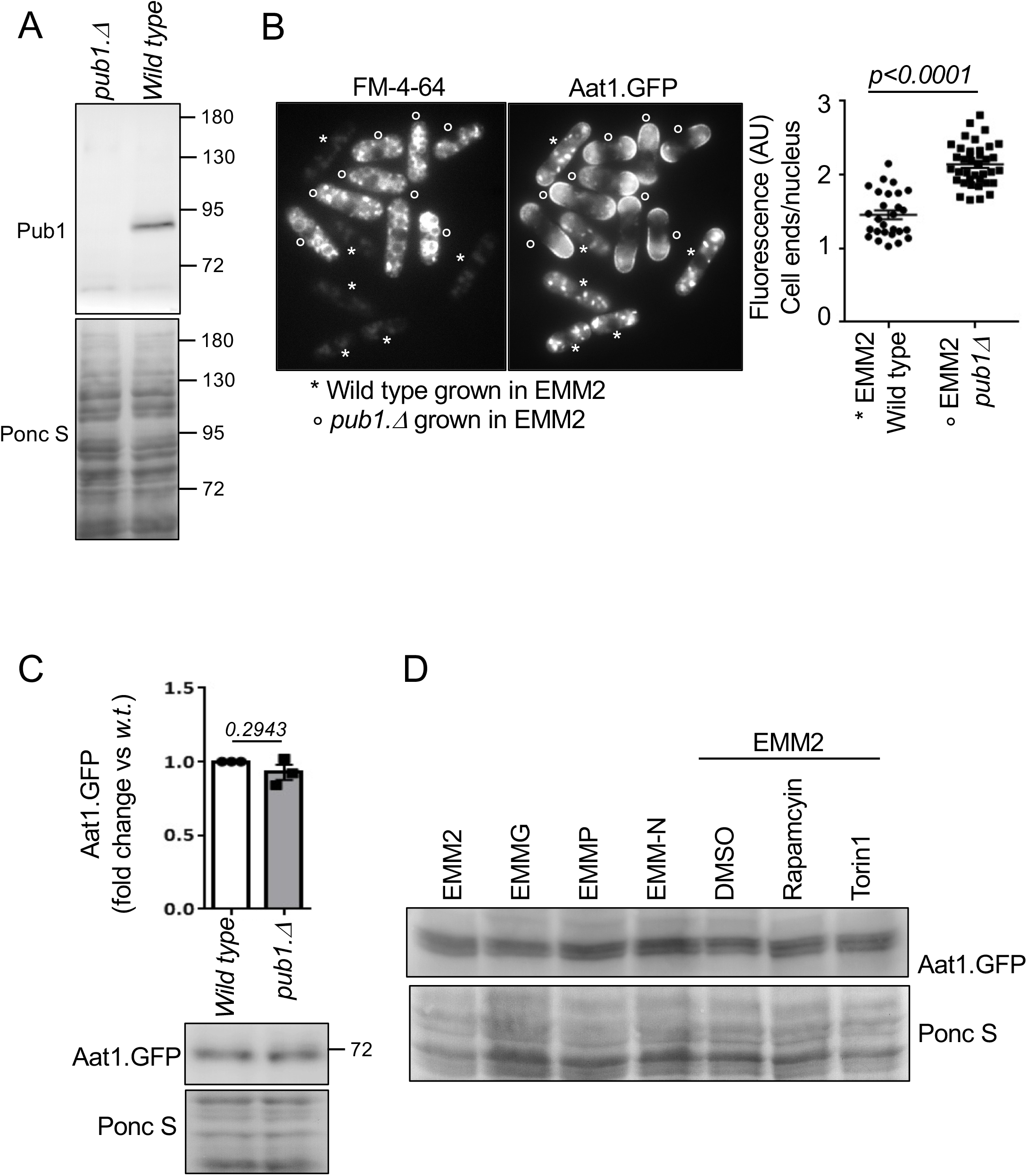
Aat1.GFP levels are unaffected by Pub1 and growth media. A) anti-Pub1 recognise Pub1. Protein extracts were prepared from indicated cells and immunoblotted for Pub1 Ponceau S is used to stain total protein. B) FM-4-64 does not interfere with Aat1.GFP fluorescence (compare with figure 2A). Live cell imaging of Aat1.GFP in indicated strains. To differentiate between the two cell types (which were mixed at a ratio 1:1 immediately prior to imaging) *pub1.Δ* cells were initially stained with FM-4-64 (indicated by a circle) for 45 min (accumulates in the vacuoles), before they were combined with unstained cells for immediate imaging. C,D) Protein extracts were prepared from indicated yeast strain or treatments and immunoblotted for GFP to visualize Aat1.GFP, Ponceau S is used to stain total protein. C) A Bars mean fold change in levels vs. EMM2 ± s.e.m., *n* = 3. *n* represent biological independent experiments. Representative immunoblots are shown.

**Supplementary Figure 2.**
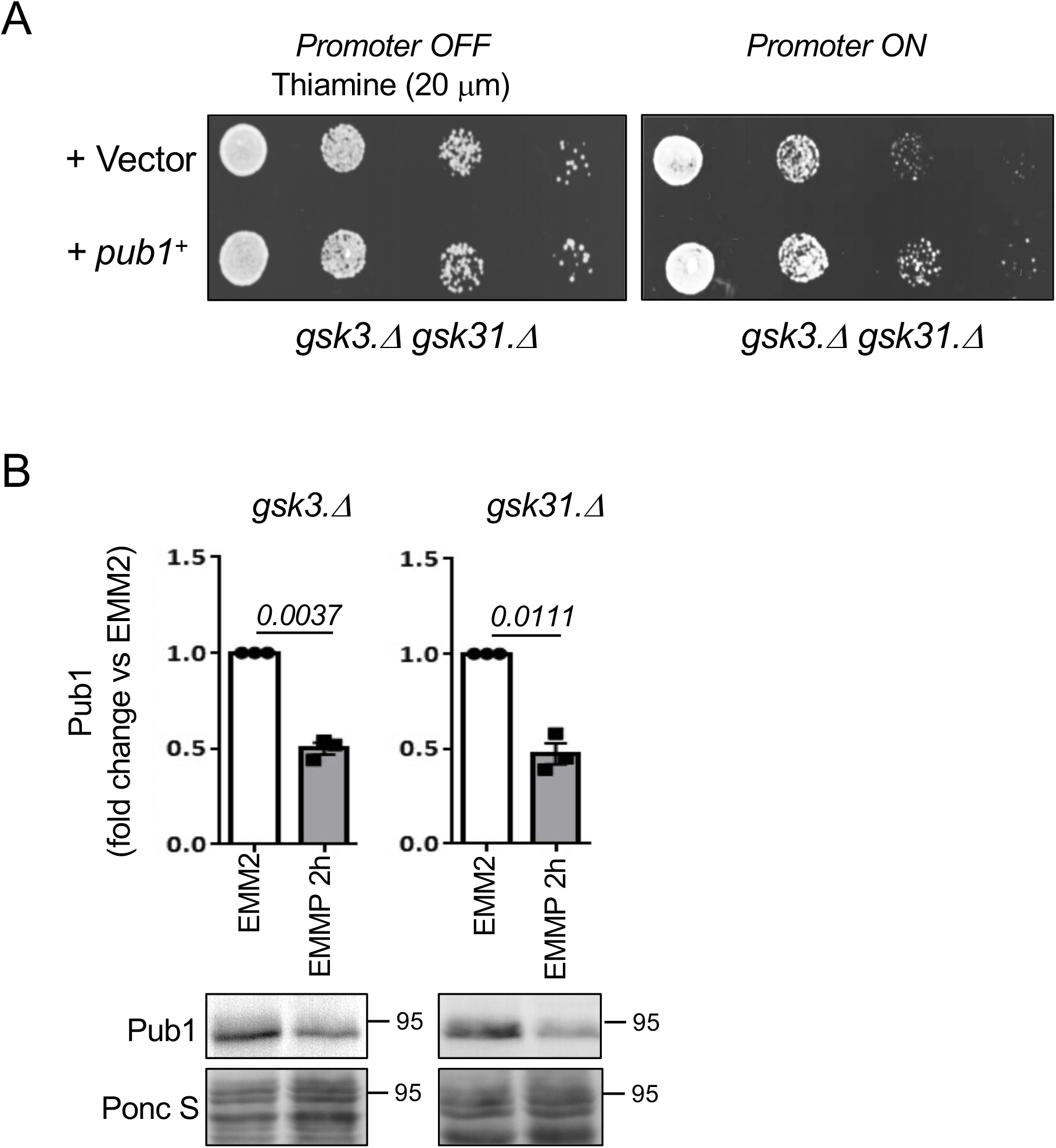
Overexpression of Pub1 cannot rescue the growth defect of cell lacking Gsk3 activity. A) Growth characteristics of indicated strains on EMM2 and EMM2 + 20 μg/ml thiamine to repress promoter. similar results were obtained for three independent biological repeats. B) Protein extracts were prepared from indicated yeast strains and treatments and immunoblotted for Pub1, Ponceau S is used to stain total protein. Bars indicate fold change in levels vs. EMM2 ± s.e.m., *n* = 3. *n* represent biological independent experiments. Representative immunoblots are shown.

**Supplementary Figure 3.**
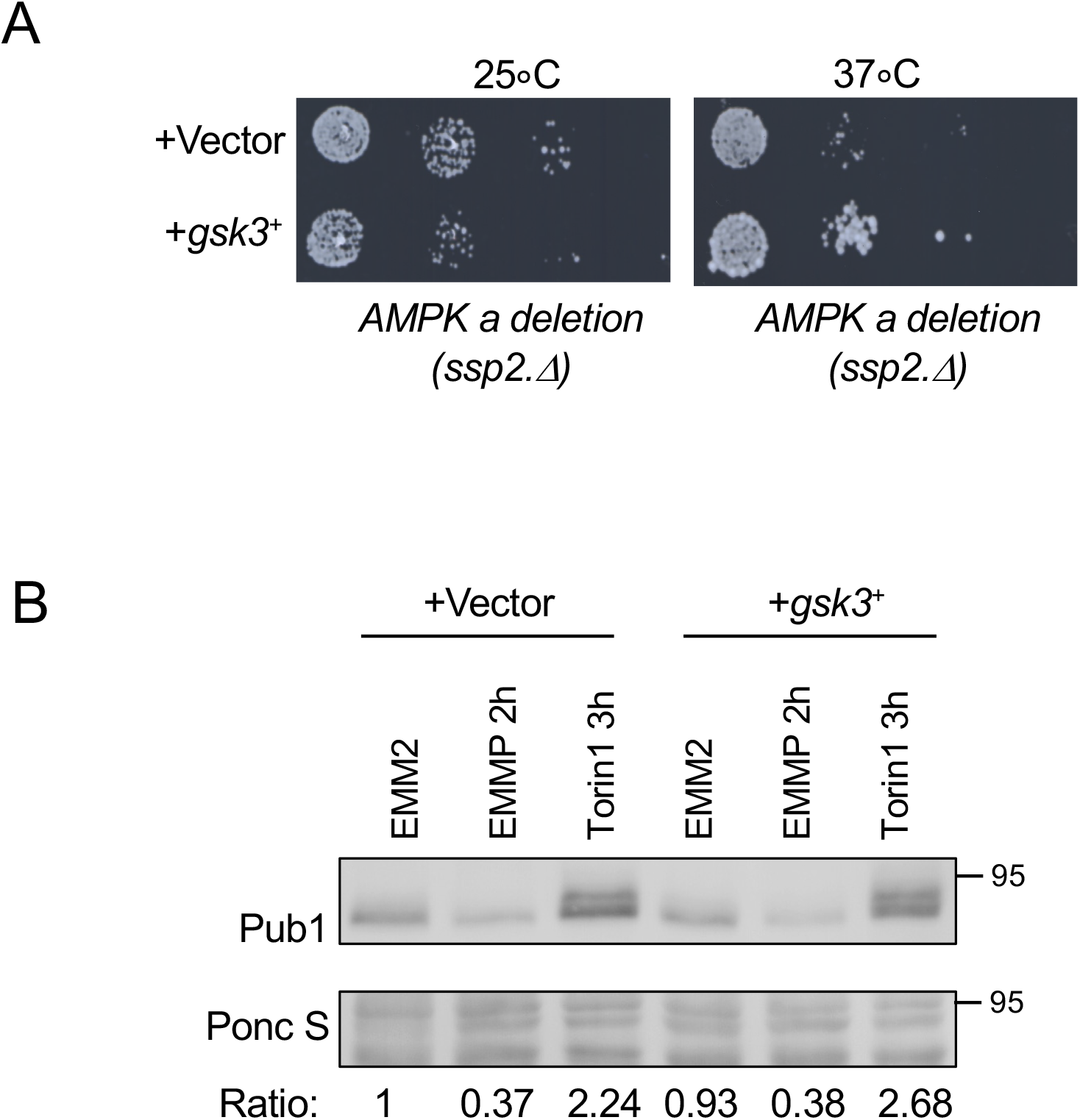
Over-expression of Gsk3 in wild type cells do not block Pub1 regulation under nitrogen-stress. A) Over-expression of Gsk3 rescue growth of cell lacking AMPK activity. Growth characteristics of indicated strains on EMM2 at 25 and 37 degree. Similar results were obtained for three independent biological repeats. B) Protein extracts were prepared from indicated yeast strains and treatments and immunoblotted for Pub1, Ponceau S is used to stain total protein. Ratio’s indicate fold change in Pub1 levels vs EMM2.

**Supplementary Figure 4.**
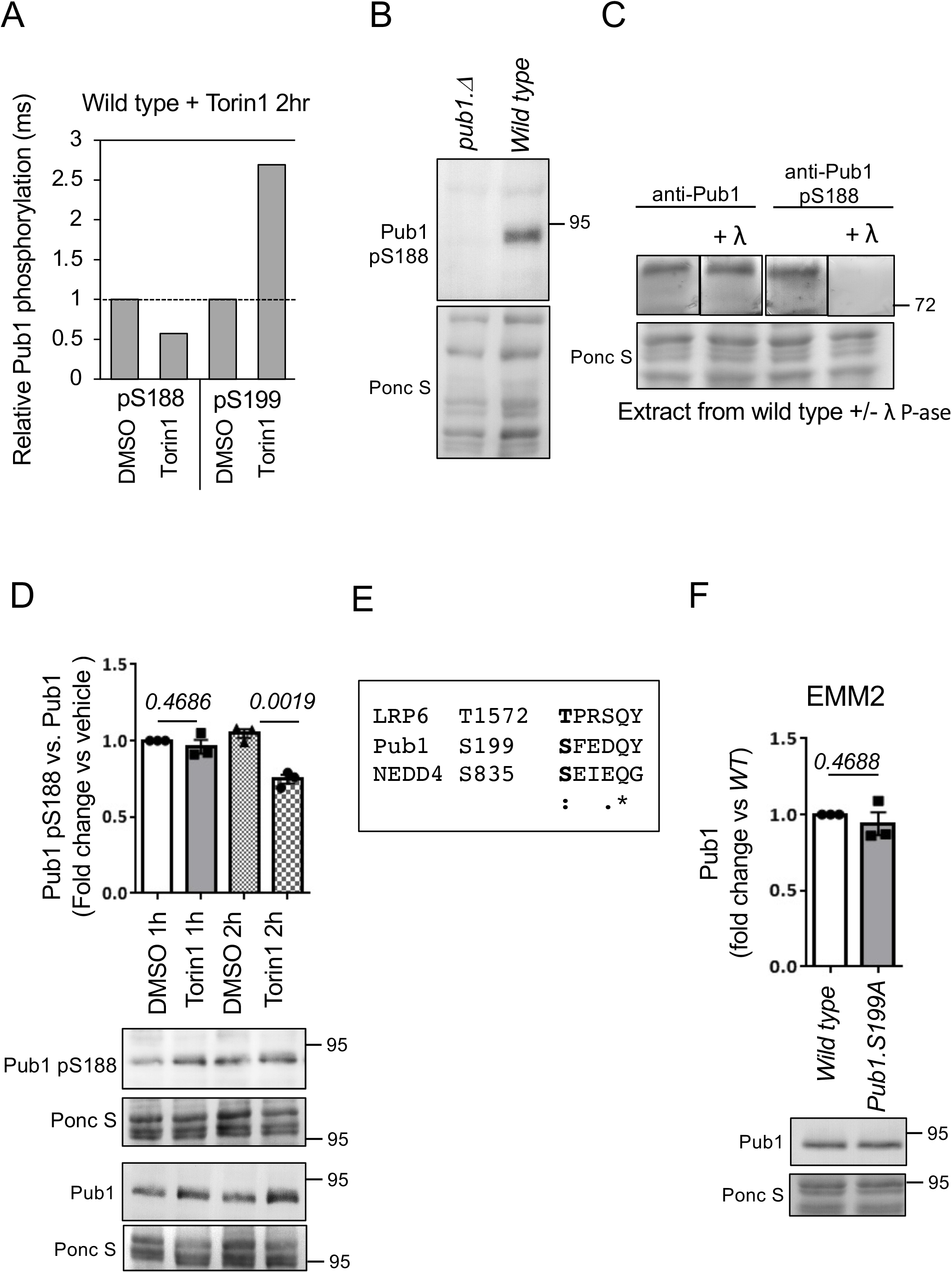
**A)** Chart illustrating changes to phosphorylation of Pub1 S188 and S199 following Torin1 treatment for 2hr by mass spectrometry (see supplementary Table 1). B-D, F) Protein extracts were prepared from indicated yeast strains and treatments and immunoblotted for Pub1 or Pub1 pS188, Ponceau S is used to stain total protein. B, C) anti-Pub1 S188 recognise Pub1 and is phospho-specific. C) Indicated Ponceau S stained PVDF membranes were cut and incubated with or without lambda phosphatase for 1 hour, to dephosphorylate proteins prior to immuno-blotting. D) 2 hours of Torin1 treatment reduce phosphorylation of Pub1 serine 188. F) Pub1 levels are unaffected by the status of serine 199 phosphorylation in good nutrient environments. E) sequence alignment of LRP6 (an established GSK3 substrate, which do not need a priming kinase) Pub1 S199 and human NEDD4 S835, also see sequence alignment in figure 6A.

**Supplementary Table 1.**
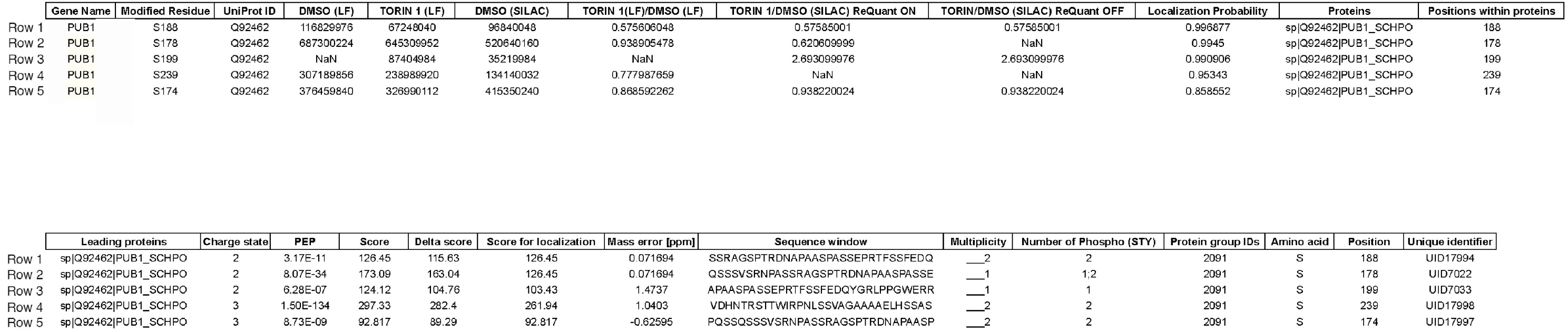
Results from mass spec analysis of SILAC labelled cells treated with Torin1 for 2 hours.

